# Detection of PD-L1-expressing myeloid cell clusters in the hyaluronan-enriched stroma in tumor tissue and tumor-draining lymph nodes

**DOI:** 10.1101/2020.12.15.422923

**Authors:** Paul R. Dominguez-Gutierrez, Elizabeth Kwenda, William Donelan, Padraic O’Malley, Paul L. Crispen, Sergei Kusmartsev

**Affiliations:** Department of Urology, University of Florida, Gainesville, Florida, USA

## Abstract

Expression of the transmembrane protein PD-L1 is frequently up-regulated in cancer. Since PD-L1-expressing cells can induce apoptosis or anergy of T lymphocytes through binding to PD1 receptor, the PD-L1-mediated inhibition of activated PD1^+^ T cells considered as a major pathway for tumor immune escape. However, the mechanisms that regulate the expression of PD-L1 in the tumor microenvironment not fully understood. Analysis of organotypic tumor tissue slice cultures, obtained from tumor-bearing mice as well as from cancer patients, revealed that tumor-associated hyaluronan (HA) supports the development of the immunosuppressive PD-L1^+^ macrophages. Using genetically modified tumor cells, we identified both epithelial tumor cells and cancer-associated fibroblasts (CAFs) as the major source of HA in the tumor microenvironment. HA-producing tumor cells and, in particular CAFs of bone marrow origin, directly interact with tumor-recruited Hyal2^+^ myeloid cells forming the large stromal congregates/clusters that are highly enriched for both HA and PD-L1. Furthermore, similar cell clusters comprising of HA-producing fibroblasts and PD-L1^+^ macrophages were detected in the tumor-draining lymph nodes. Collectively, our findings indicate that the formation of multiple large HA-enriched stromal clusters that support the development of PD-L1-expressing antigen-presenting cells in the tumor microenvironment and draining lymph nodes could contribute to the immune escape and resistance to immunotherapy in cancer.

## Introduction

The immunosuppressive ligand PD-L1 plays an important role in the regulation of T cell-mediated immune response and tumor-associated immune tolerance (**1**). Recent studies demonstrated that PD-L1 expression by host’s myeloid APCs, is essential for PD-L1 mediated immune evasion and immunotherapy (**2–4**). Bladder cancer is characterized by a highly immunosuppressive microenvironment including up-regulated expression of inhibitory ligand PD-L1 (**5, 6**) and the strong presence of immunosuppressive myeloid cells such as myeloid-derived suppressor cells (MDSCs) and tumor-associated macrophages (TAMs) (**7–9**). The tumor cells can up-regulate the PD-L1 expression in tumor-associated macrophages (**6**), however, the mechanism of tumor-mediated regulation of PD-L1 expression in myeloid cells remains unresolved.

Tumor stroma plays a major role in tumor growth and is comprised of both cellular and extracellular components. Bladder cancer is enriched for both cancer-associated fibroblasts (CAFs) as well as for one of the major components of extracellular matrix hyaluronan (HA) **(10–11**). CAFs play very diverse roles in tumor development and progression including stimulation of tumor-promoting inflammation, tumor proliferation, and invasion, neovascularization as well as tumor-associated immune suppression (**12–14**). Several lines of evidence suggest a strong interplay between CAFs and myeloid cells. Thus, fibroblasts play pivotal roles in the recruitment of CCR2-expressing monocytic myeloid cells and polarization of those recruited cells to the M2 macrophages or immunosuppressive MDSCs (**15, 16**). Roles of CAFs in the recruitment of monocytes/macrophages to the tumor tissue supported by fibroblast’s production of monocyte-macrophage chemotactic factor CCL2. Moreover, HA deficiency in tumor stroma resulted in a marked reduction of both macrophage recruitment and tumor neovascularization (**16–17**).

Here we demonstrate that tumors stimulate the gathering of stromal cells into large cell clusters. The major cellular components of these clusters include HA-producing fibroblasts, tumor epithelial cells, and macrophages. Similar stromal structures with HA-producing fibroblasts and PD-L1^+^ macrophages were detected in tumor-draining, but not in distant lymph nodes. Our results support the idea that stroma-produced HA directly supports the development of immunosuppressive PD-L1-expressing macrophages

## Results

### Tumor stromal cells are enriched for HA and associated with large cell conglomerates of PD-L1-expressing macrophages

To explore the development of PD-L1^+^ macrophages in the tumor microenvironment, we utilized the organoid tumor tissue-slice technique. To this end, we injected MBT2, the murine bladder tumor cells into syngeneic C3/He mice surgically excised the developed tumors, and prepared the precision-cut 200-300 micron tumor tissue slices using the Compresstome tissue slicer. Prepared tissue-slices are cultured in 24-well plates in a complete culture medium. The viability of tissue slices was monitored using the Live/Dead viability assay (Invitrogen).

Six to twelve hours after the initiation of tumor tissue slice cultures, we noticed the development of multiple stromal cell conglomerates/clusters firmly attached to the plastic (**Fig. 1a and Supporting Fig. S1**). Staining for PD-L1 revealed that these cell clusters are highly positive for this marker. Intriguingly, the stromal cell clusters are highly enriched for the HA. The majority of the PD-L1^+^ cells were also positive for pan-hematopoietic marker CD45 (**Supporting Fig. S2a**). These data implicate the potential involvement of tumor stroma-associated HA in the development of PD-L1^+^ cells. A recently published study indicates that lymph nodes) could be subjected to the preparation of tissue slices to study *ex vivo* immune response (**18**). Accordingly, we collected from MBT2 tumor-bearing mice the tumor-draining and distant control LNs. Freshly collected LNs were used for the preparation of precision-cut tissue slices. A few days later, the LN tissue slice cultures were stained for the presence of PD-L1 and HA. To our surprise, the draining LNs (**Fig.1b)**, similar to the tumor tissue slices, was also able to develop the HA-enriched adherent stromal cell clusters with incorporated PD-L1^+^ macrophages. In contrast, the distant LNs were not able to produce any HA-enriched PD-L1^+^ stromal cell clusters that we observed in the cultures prepared from tumor tissues or tumor-draining LNs. Co-staining the tumor stroma with antibodies against PD-L1, HA, and macrophage marker F4/80 revealed (**Supporting Figs. S2b and S3a**) that cluster-associated PD-L1^+^ cells co-express F4/80. In addition to the mouse bladder tumor model, similar stromal HA-enriched PD-L1 expressing clusters were observed in clinical cancer tissue samples obtained from patients with bladder cancer (**Fig.2a and Supporting Fig. S3b**). Taken together, these data indicate that tumor stroma and stroma-associated HA support the development of PD-L1^+^ macrophages.

**Figure 1.**
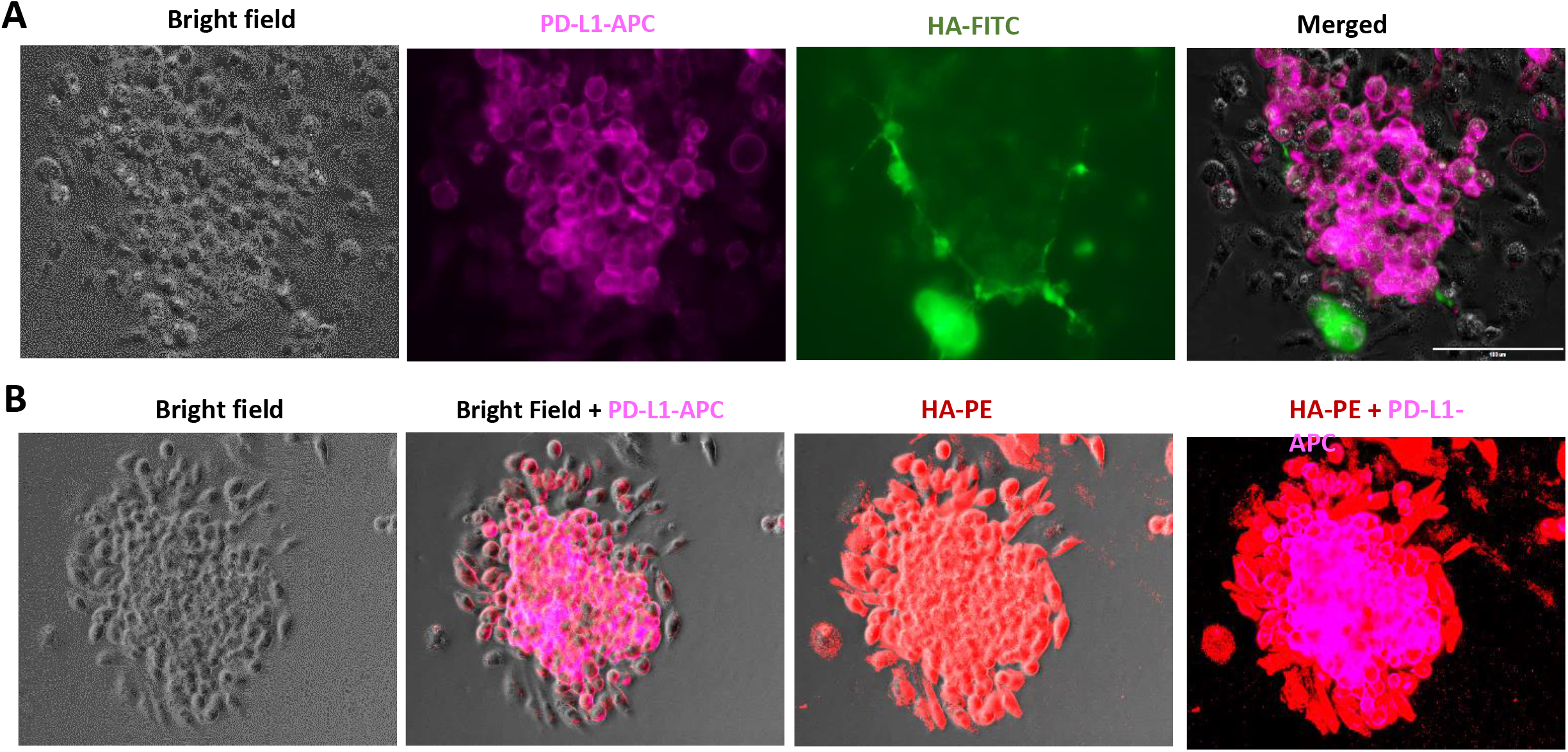
Stroma-associated HA supports the development of PD-L1^+^ cells in tumor tissue and tumor-draining lymph nodes. MBT2 tumor cells were injected into C3/He mice. Two weeks later mice were sacrificed, and tumors were surgically excised. Tissue slices prepared using tumor tissue pieces (**A**) or tumor-draining lymph node (**B**) were cultured in a complete culture medium. Plates were fixed with 4% formaldehyde and stained for the expression of PD-L1 and HA. Representative images of tumor stroma are shown. Scale bar 100 μm.

**Figure 2.**
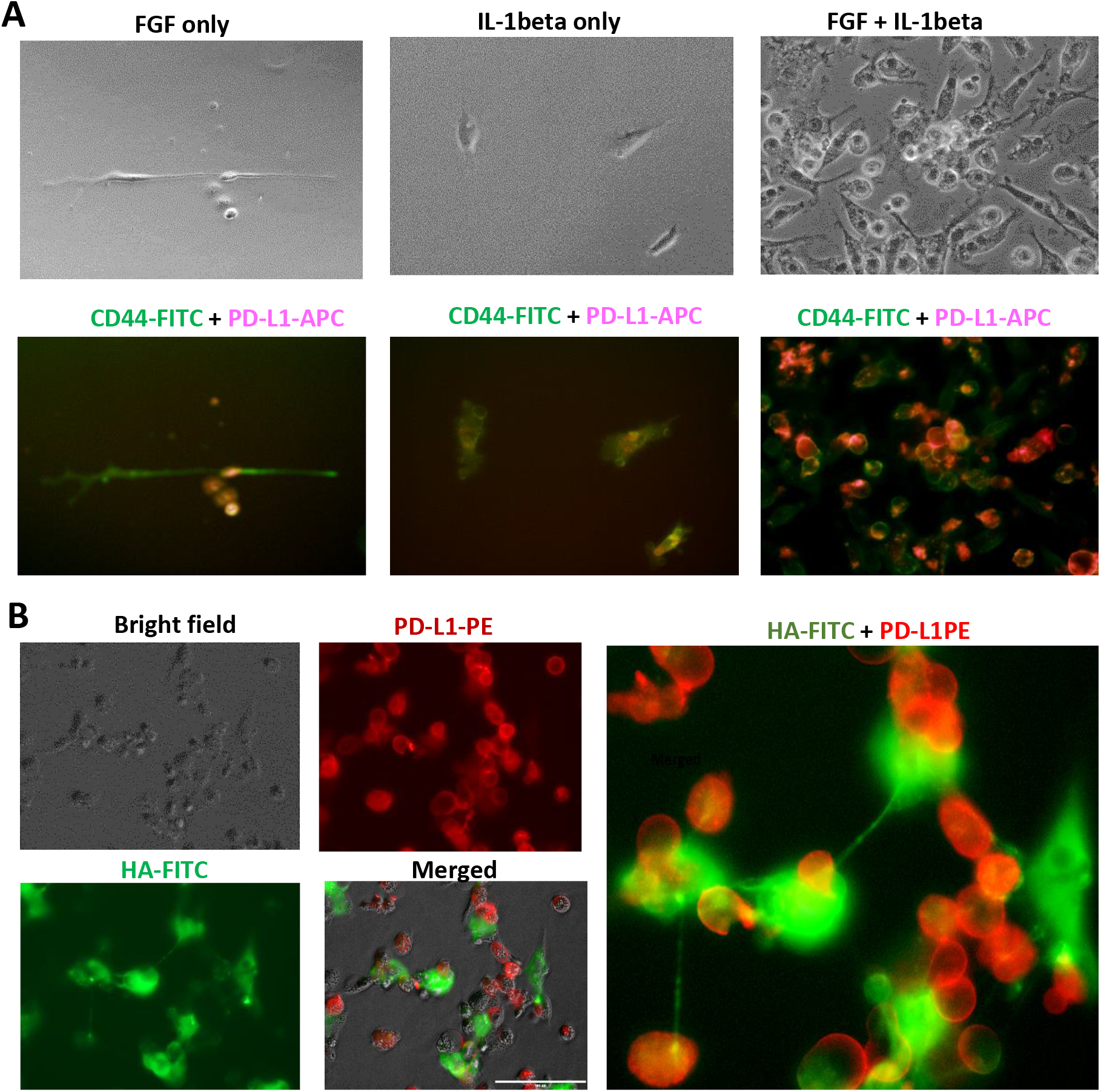
IL-1β and FGF2 promote formation of stromal PD-L1 clusters in bone marrow cell cultures. **A:** Naïve bone marrow cells cultured in 24-well plate in the presence of recombinant murine IL-1β and basic FGF. On day seven, expression of PD-L1 (red) and CD44 (green) was evaluated using IF microscopy. Representative images are shown. **B**: HA produced by stromal fibroblasts directly interacts with PD-L1^+^ macrophages. MBT2 tumor tissue slices were cultured for 24 hours. Fixed cultures stained to evaluate the expression of PD-L1 (red) and HA (green). Representative images are shown.

### HA-producing fibroblasts of bone marrow origin contribute to the development of PD-L1^+^ macrophages

Cancer-associated fibroblasts (CAFs) represent one of the major components of tumor stroma. CAFs exert diverse functions, including matrix deposition and remodeling, extensive reciprocal signaling interactions with tumor cells and infiltrating immune cells (**10**). The origin of these cells not fully understood, however some CAFs clearly demonstrate a bone marrow origin (**19**). In addition, it was reported that bone marrow-derived mesenchymal cells with fibroblast-like appearance constitutively produce HA (**20**). Here we show that bone marrow-derived cells obtained from normal naïve mice and stimulated with recombinant fibroblast growth factor (FGF2) and IL-1β give a rise to HA-producing fibroblast-like cells and PD-L1^+^ macrophages in the absence of tumor cells (**Fig.2a**). It appears that IL-1β and FGF2 have a synergistic effect on development of HA-enriched PD-L1^+^ clusters, because stimulation of bone marrow cells separately with these cytokines results in much weaker effects. Furthermore, similar clusters comprising of PD-L1^+^ macrophages and fibroblast-like HA-producing cell also detected in tumor-draining lymph nodes (**Supporting Fig. S4a**). Collectively, these data suggest that HA-producing fibroblasts of bone marrow origin could contribute to the development of PD-L1^+^ macrophages in tumor-free conditions.

To get a better insight into the formation of stromal cell clusters in tumor microenvironment, we next applied the time-lapse video take during consecutive 96 hours after initiation of tumor-slice culture. Data presented in **Supporting Figs S4b, S5** and tame-lapse video (not shown) demonstrate that formation of macrophage-fibroblast clusters is highly dynamic and interactive process, when both large irregularly-shaped fibroblast-like cells and smaller round-shaped macrophages moving around each other during formation of stromal cells clusters. **Fig.2b** illustrates that fibroblast-secreted HA (green) physically interacts with PD-L1^+^ macrophages (red). These data clearly indicate that HA is directly involved in the development of PD-L1^+^ macrophages.

### Both fibroblasts and tumor cells contribute to the development of HA-enriched PD-L1-expressing cell clusters

To delineate the roles of CAFS and epithelial tumor cells in the development of PD-L1^+^ cells, we stably transfected murine MBT2 tumor cell line with GFP using lentivirus. GFP-expressing tumor cells were injected in mice, and 2 weeks later harvested tumor was used for the preparation of tumor tissue-slices (**Fig.3a**). Data presented in **Fig. 3b** demonstrate that initially PD-L1-positive clusters (magenta) and GFP-positive epithelial tumor cells (green, central panel) are frequently localized separately. PD-L1^+^ cells are associated mostly with GFP-negative fibroblast-like cells. However, a few days later when tumor stroma becomes denser, the PD-L1^+^ cells, fibroblasts, and tumor cells all become mixed in stromal cell clusters (**Supporting Fig. S6a**). Our data also demonstrate that both tumor cells and CAFs produce HA. In addition to the bladder tumor model, we also observed the development of PD-L1-expressing cell clusters in murine colorectal CT26-GFP^+^ tumors (**Supporting Fig. S6b**). Similar to MBT2-GFP tumor model, tumor tissue-slice cultures from CT26-GFP tumor-bearing mice produced the HA-enriched tumor stroma (red) comprised of GPF-positive epithelial tumor cells (green) and PD-L1^+^ round-shaped macrophages (magenta). Taken together, our data indicate that HA-producing fibroblasts play initiating and supporting roles for the development of PD-L1^+^ macrophages, while epithelial HA-producing tumor cells became incorporated into stromal macrophage-fibroblast clusters later.

**Figure 3.**
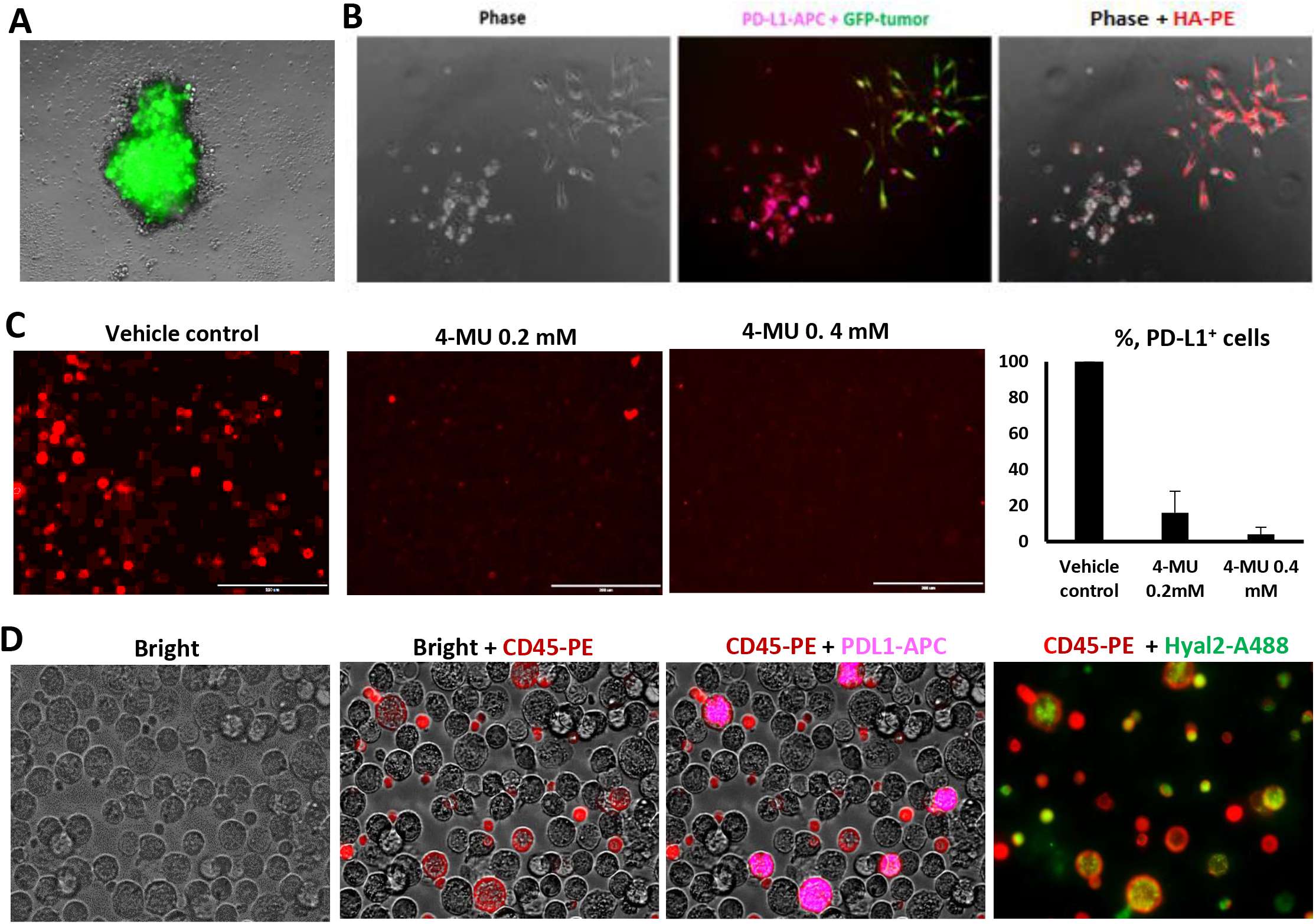
Identification of HA-producing cells in tumor stroma. GFP-expressing MBT-2 tumor cells were injected in mice, and two weeks later the harvested tumor was used for the preparation of tumor tissue slice culture. Tumor tissue slices were cultured for 72 hours, then fixed and stained with PD-L1-APC and HA-PE Abs. Representative images of whole GFP^+^ tumor piece (**A**) or tumor stroma (**B**) are shown. **C:** Gr-1^+^ cells were enriched using magnetic beads from the spleen of MBT2 bearing mice. Myeloid cells (6×10^5^ cells/per well) were mixed with MBT-2 tumor cells (3×10^5^ cells per/well) and added to the 24-well plate in a complete culture medium. HAS inhibitor 4-MU was added at indicated concentrations immediately after the plating of myeloid and tumor cells. On Day 7 cells were collected and stained with anti-PD-L1-PE Abs. Average means ± SD shown; *, P<0.05. **D:** Tumor-induced PD-L1^+^ myeloid cells co-express Hyal2. Gr-1^+^ cells were isolated from MBT2 bearing mice. Myeloid cells were mixed with MBT-2 tumor cells and cultured in 24-well plates. On day 5 cells were collected and stained with antibodies against CD45 (red), PD-L1 (magenta), and Hyal2 (green). Representative images are shown.

### Tumor cells promote the development of PD-L1^+^ macrophages in an HA-dependent manner

The co-culture of the tumor cells with Gr-1^+^ MDSCs leads to the up-regulation of PD-L1 expression and development, promoting differentiation of myeloid-derived cells into PD-L1^+^F4/80^+^ macrophages (Ref. 6 and **Supporting Fig. S7a)**. MBT2 tumor cells produce HA (**Fig. 3b**) because of high expression of hyaluronan synthase 3 (HAS3) (**Supporting Fig. S7b**). To examine the potential involvement of tumor-produced HA in the tumor-induced up-regulation of PD-L1 expression in myeloid cells, we have co-cultured MBT2 tumor cells and murine Gr-1^+^ cells with added HAS inhibitor 4-methylumbelliferone (4-MU). Data presented in **Fig. 3c** demonstrate that inhibition of HA synthesis results in a dose-dependent reduction of PD-L1 expression.

The MDSCs may affect the HA metabolism in tumor tissue through membrane-bound enzyme hyaluronidase 2 (Hyal2) (**21**). Hyal2 is a rate-limiting enzyme, which upon activation able to degrade the extracellular HA into small fragments with low molecular weight (LMW-HA), and its expression was increased in both tumor-associated and blood-derived myeloid cells in cancer patients with bladder cancer. To examine whether Hyal2-expressing cells could potentially contribute to the HA-mediated development of PD-L1^+^ macrophages, we co-cultured Gr-1^+^ MDSCs and MBT2 tumor cells for 5 days and then co-stained with CD45 (pan-hematopoietic marker), PD-L1, and Hyal2. Data presented in **Fig.3d** indicate that PD-L1^+^ cells co-express the Hyal2 enzyme. Next, we checked the tumor-associated CD11b myeloid cells **(Fig.4a**) for the expression of PD-L1 and Hyal2. To this end, we isolated CD11b cells from murine MBT2 bladder tumor tissue and stained for those markers. Data presented in **Fig. 4b and Supporting Fig. S8a** indicate that majority of PD-L1^+^ cells also-co-express the Hyal2 enzyme.

**Figure 4.**
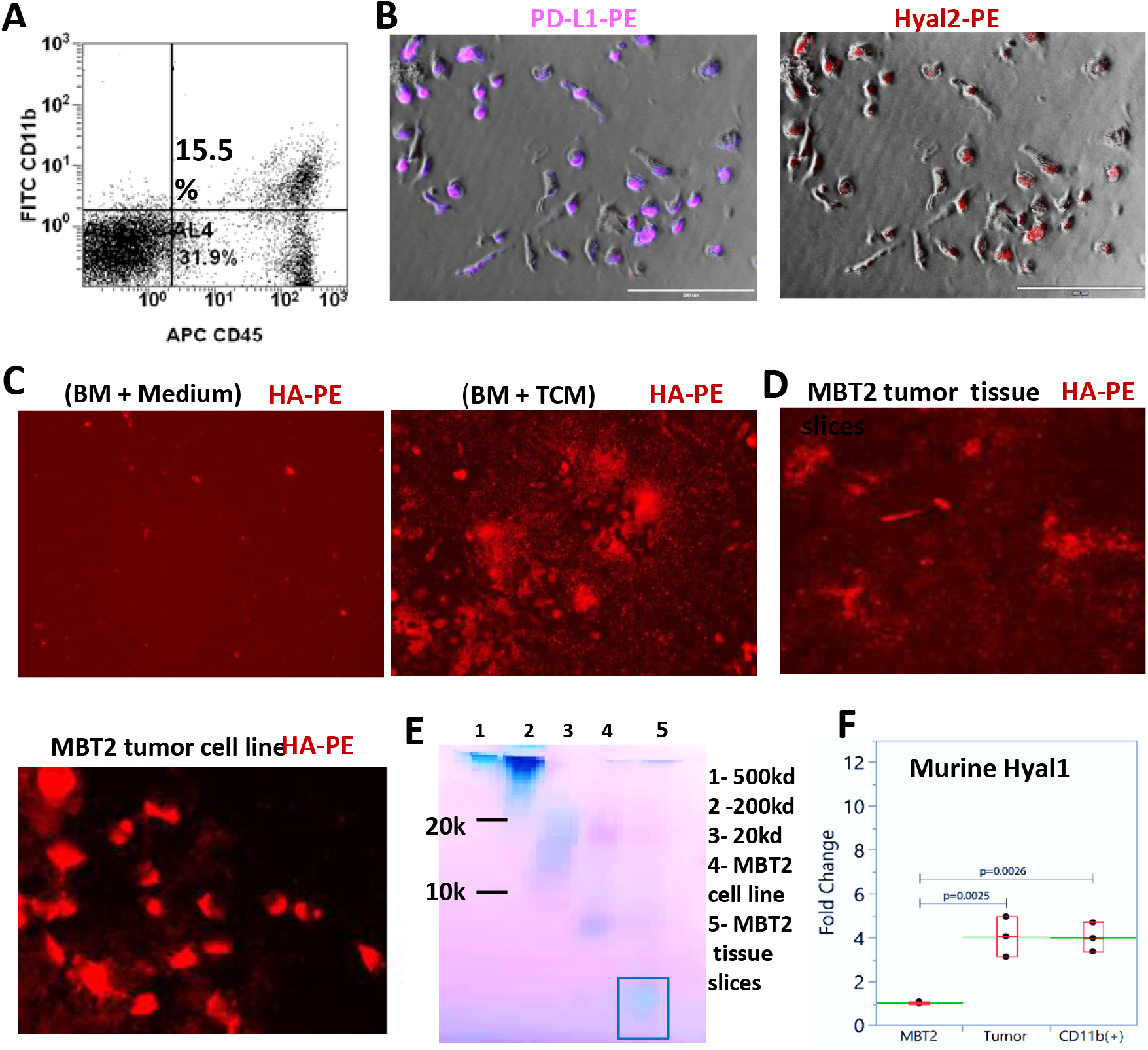
TCM stimulates expression of membrane-linked Hyal2 in bone-marrow-derived myeloid cells and promotes degradation of extracellular HA. **A:** Murine bladder tumor tissue infiltrated with CD11b myeloid cells. Tumors surgically resected from MBT2 tumor-bearing mice were digested with collagenase cocktail to prepare single-cell tumor suspension. The single-cell suspension was co-stained with CD11b-FITC and CD45-APC antibodies and analyzed by flow cytometry. The results of one representative experiment are shown. **B:** Tumor-associated CD11b myeloid cells express Hyal2 and PD-L1. Tumor-infiltrating myeloid cells from MBT2 tumors were purified with magnetic anti-CD11b beads. Isolated cells were placed in 24 well plates and cultured overnight allowing cells to adhere to the plate. The next day cells were fixed and stained for and PD-L1 (magenta, left panel). And Hyal-2 (red, right panel), **C**: Exposure of BM-derived myeloid cells to TCM stimulates the fragmentation of extracellular HA. 24-well plates were pre-coated with sterile commercial HA (200 kDa). CD11b myeloid cells isolated from murine naïve BM were cultured in the presence or absence of TCM. On day 7 cell cultures were fixed with 4% formaldehyde and stained for HA. HA expression (red) was examined by the IF microscope. Representative images are shown. Scale bar 200 μm. **D**: Visualization of murine tumor-produced HA. The MBT2 tumor tissue slices (left image) or MBT22 tumor cell line (right image) alone were cultured for 72 hours. Tumor cells were carefully removed using an enzyme-free approach, and tumor-free wells were stained for tumor-produced HA. To visualize tumor-produced HA, plate wells were subsequently incubated with biotinylated HA-binding protein and PE-labeled Streptavidin. Images were obtained using IF microscopy. **E:** Electrophoretic analysis of HA produced by the MBT2 tumor cell line and MBT2 tumor tissue slices. Cell-free supernatants collected from the tumor cell line and tumor tissue slice cultures were treated with ethanol, proteinase K, and benzonase before applying samples to polyacrylamide electrophoresis. Commercial HA with MW 20, 200, and 500 kDa were used as control. Hyaluronan was visualized by staining with “Stains All” dye. **F:** Expression of Hyal1 mRNA in MBT2 tumor and tumor-infiltrating myeloid cells. Expression of Hyaluronidase 1 (*Hyal1*) in MBT2 bladder tumor cell line and MBT2 tumor-bearing mice. Total RNA from MBT2 tumor cells (MBT2), whole tumor resected from MBT2 tumor-bearing mice (tumor), and tumor-infiltrating CD11b myeloid cells were extracted using Trizol reagent. The expression of HyaL1 mRNA was measured using qRT-PCR. Average means ± SD shown; n=3; *, P<0.05.

The activity of Hyal2 in myeloid cells can be stimulated with tumor-conditioned medium (TCM) or IL-1β (**21**). To examine whether this activation is associated with the up-regulation of PD-L1 expression, we used the murine CD11b cells isolated from normal bone marrow. Data presented in **Fig.4c** demonstrate that similarly to its’s human counterpart, the TCM-activated murine myeloid cells able to degrade extracellular HA. A similar extent of HA degradation was detected while analyzing HA produced by MBT2 tumor-tissue slices, but not by the MBT2 tumor cell line (**Fig.4d**). These data were confirmed by gel electrophoresis (**Fig. 4e**). Specifically, the electrophoretic analysis confirmed that HA produced by MBT2 tumor cell line consisted of fragments with intermediate size (MW ~20 kDa), whereas tumor tissue slices-derived HA showed lower molecular weight (<10 kDa). Since the Hyal2 enzyme specifically degrades HA to fragments with MW 20 kD (**22, 23**), other types of hyaluronidases could likely be involved.

To address this question, we measured the expression of Hyal1 and Hyal3 using qRT-PCR in MBT2 tumor cell line, whole tumor tissue from tumor-bearing mice, and CD11b cells isolated from the tumor. Levels of Hyal3 were very low, whereas the expression of Hyal1 was up-regulated in myeloid cells as compared to the MBT2 tumor cell line (**Fig.4f**). Hyal1 is an intracellular lysosomal enzyme that degrades the internalized HA to very small fragments with low molecular weight less than 5kD (LMW-HA). It has been proposed that membrane-bound enzyme Hyal2, which breaks down the extracellular HA, works in concert with Hyl1 to produce the LMW-HA (**22–23**). Taken together, our data indicate that is very likely that both the Hyal2 and Hyal1 enzymes are involved in the degradation of tumor-associated HA.

## Discussion

A better understanding of stroma-immune interactions in the tumor microenvironment could shed a light on the roles of tumor stroma in the regulation of anti-tumor immune response. However, most of the classical research approaches to study the tumor microenvironment have significant limitations. One of the popular methods is a preparation of single tumor suspension using mechanical and enzymatic digestion of tumor tissue. The resultant single tumor cell suspension is suitable for the analysis of immune tumor-infiltrating cells using flow cytometry or immune fluorescence, isolation of certain cell subsets, etc. However, since mechanical and enzymatic tissue digestion leads to the disruption of naturally occurring cell-cell interactions, this approach does not work for studies focused on tumor stroma and stroma-immune interaction. Another research method to study the tumor tissue is immunohistochemistry (IHC) of formalin-fixed or frozen tissues. This method works in many cases, however, because of the high complexity of the tumor microenvironment, it is very challenging to study the mechanisms of stroma-immune interactions using IHC.

In contrast, the cultures of precision-cut tissue slices prepared from freshly excised experimental or clinical tumors create nearly ideal conditions to explore the interaction between tumor stroma and immune cells. Once tumor slices are placed in a culture flask or plate, they start the formation of adherent stroma that includes an extracellular matrix with attached fibroblasts, macrophages, and other immune cells. Using the GFP-expressing tumor models, we noticed that initial stromal clusters were tumor cell-free and comprised mainly of HA-producing fibroblasts and myeloid cells. However, a few days later, tumor cells eventually migrated from tissue slices and incorporated them into the stroma. Both epithelial tumor cells and cancer-associated fibroblasts were able to produce the HA which directly interacted with myeloid cells and supported the development of PD-L1^+^F4/80^+^ macrophages. These stromal cell clusters are dynamic structures that quickly grow over time in size and numbers. We also noticed that more vascularized tumors produced higher numbers of stromal clusters. Multiple cell structures with incorporated HA-producing fibroblasts and PD-L1^+^ round-shaped myeloid cells were detected in several tumor types including bladder carcinoma MBT2, murine colon carcinoma CT26 **(Supporting Fig. 6b)** and kidney carcinoma Renca (**Supporting Fig. S8b**). These findings indicate that the formation of multiple stromal clusters enriched for HA-producing fibrobalasts, tumor cells and PD-L1^+^ antigen-presenting cells may represent a general mechanism of immune escape in cancer.

HA has been implicated in regulating a variety of cellular functions in both tumor cells and tumor-associated stromal cells, suggesting that altered HA levels can influence tumor growth and malignancy at multiple levels. Previously published studies demonstrate that HA increases the proliferation rate of tumor cells *in vitro* and promotes cell survival under anchorage-independent conditions (**11, 26)**. At the molecular level, HA activates the PI3K/Akt pathway and influences the expression of cell cycle regulators. Furthermore, HA also can inhibit tumor cell apoptosis, as demonstrated by experimentally modifying HA levels. Importantly, increased HA production in cancer is frequently associated with enhanced HA degradation due to high levels of expression/activity of hyaluronidases (**27, 28**). Increased HA degradation leads to the accumulation of HA fragments with low molecular weight (LMW-HA) (**29–30**). LMW-HA seems to have specific pro-tumoral functions by promoting inflammation, tumor angiogenesis, and metastasis through stimulating the production of cytokines, chemokines, growth factors in TLR2/TLR4 dependent manner (**31).** Conversely, the high molecular weight HA shows anti-inflammatory and anti-oncogenic effects **(32, 33**). Our data support the idea that in addition to cancer-related inflammation and tumor angiogenesis, the tumor-associate HA also involved in the immune suppression in cancer, since HA supports the development of immunosuppressive PD-L1^+^ macrophages in both tumor tissues and TDLNs.

Tumor-draining lymph nodes (TDLNs) are essential for the initiation of an effective antitumor T-cell immune response. However, tumors may affect the immune-initiating function of TDLNs (**24**). A recently published study demonstrated that TDLNs in tumor-bearing mice are enriched for both PD-L1^+^ antigen-presenting cells and tumor-specific PD-1^+^ T cells (**25**). TDLN-targeted PD-L1-blockade induced enhanced anti-tumor T cell immunity by seeding the tumor site with T cells, resulting in improved tumor control. Moreover, abundant PD-1/PD-L1-interactions in TDLNs were also observed in non-metastatic melanoma patients. Our data demonstrate that similarly to the tumor tissue, PD-L1^+^ antigen-presenting cells in the TDLNs are associated with HA-producing fibroblasts (**Fig. 1b, Supporting Fig.4a**) and characterized by enhanced degradation of TDLN-associated HA (**Supporting Fig. S9**). Taking into account a prominent role of PD-L1^+^ cells in immune tolerance and tumor-associated immunosuppression, it is plausible that observed cell clusters comprising of HA-producing cells and PD-L1^+^ antigen-presenting cells could contribute to the formation of the immunosuppressive and immune tolerogenic microenvironment in both tumor tissue and TDLNs.

## Materials and Methods

### Reagents and culture medium

Murine recombinant FGF2 and IL-1β were acquired from R&D Systems (Minneapolis, MN). The proteinase K and Benzonase were purchased from Sigma-Aldrich. Hyaluronan with molecular weight 10, 20, 200, 500, 700, and 1000 kDa was obtained from Lifecore Biomedical (Chaska, MN). Biotinylated hyaluronan-binding protein (HABP) was supplied by Millipore-Sigma. Hyaluronidase 2 polyclonal antibody conjugated with biotin, or Alexa-488 obtained from Bioss Antibodies. All other antibodies used for immune fluorescence and flow cytometric analysis were acquired from Biolegend (San Diego, CA). *In vitro* experiments were conducted using RPMI-1640 medium supplemented with 20 mM HEPES, 200 U/ml penicillin, 50 μg/ml streptomycin (all from Hyclone) and 10% FBS from ATCC (Manassas, VA),

### Mice and tumor models

Female 6-8 wk old C3/He, BALB/c, and C57/BL6 mice were obtained from the Taconic. The murine bladder carcinoma cell line MBT-2 was purchased from JCRB Cell Bank (Japan), murine colon carcinoma CT26/GFP from GeneCopoiea (Rockville, MD), murine kidney carcinoma Renca and bladder tumor MB49 from American Type Culture Collection (Manassas, VA). Tumor cells were maintained at 37°C in a 5% CO2 humidified atmosphere in complete culture media. To establish subcutaneous tumors, mice were injected with 1×10^6^ tumor cells into the left flank of syngeneic mice.

### Generation of GFP-expressing MBT2 tumor cells

Lentivirus encoded GFP/Luc genes was acquired from GeneCopoiea (Rockville, MD). Murine bladder tumor MBT2 cells were cultured in RPMI-1640 medium supplemented with heat-inactivated FBS (10%), penicillin (100 U/ml), and streptomycin (100 μg/mL) at 37 °C and 5% CO2. For in vitro transduction, cells were transduced with 5 μl (2× 108 IU/μl) viral solution together with 5 μg/ml polybrene (EMD Millipore) for 5-7 days. The medium was replaced by fresh RPMI 1640 medium supplemented with puromycin every two days post-transduction. GFP/Luc-expressing stable tumor cell line was established by selection of GFP-positive cells cultured in pyromycin-supported culture medium.

### Preparation of organotypic tissue slices

The precision-cut 100-300-micron tissue slices of bladder tissues as wells as of draining lymph nodes were produced using a Compresstome tissue slicer VF-300-0Z. After cutting, tumor tissue slices were placed into 24-well cell culture plates in complete RPMI-1640 medium supplemented with 10% FBS and antibiotics and cultured at a humidified CO_2_ incubator for 1-2 weeks. Cell viability of cultured tissue slices was tested using the Live/Dead kit (Invitrogen).

### Visualization of tissue-associated HA

Tumor tissue slices were cultured for 3-10 days in 24-well cell culture plates in a humidified CO_2_ incubator at 37° C to allow for the production of HA. At the end of incubation, tissue-produced HA was found settled at the bottom of the culture plate wells. To monitor and visualize the accumulation of tissue-produced HA fragments on the plastic surface, the tissue slices and culture medium were removed at different time points. The empty wells were washed with warm PBS and fixed with 4% formaldehyde for 30 min. After fixation, plate wells were washed with PBS containing 2% FBS and incubated overnight with biotinylated HA-binding protein (3 μg/ml, Calbiochem-EMD Millipore) at 4° C (**34**). Next day, after washing the wells with PBS containing 2% FBS, streptavidin-conjugated with fluorochrome was added to the wells and incubated for 30 min at 4° C. Plates were then washed with PBS and the bottoms of the wells were visualized using EVOS (Invitrogen) or Lionheart (Biotek Instruments) immunofluorescent imaging microscopes.

### Evaluation of HA size

Analysis of HA molecular weight was done using polyacrylamide gel electrophoresis as described previously (**35)**. Briefly, conditioned medium tissue slices were centrifuged, aliquoted, and stored at −80° C. To prepare samples for HA size analysis, thawed samples were digested with proteinase K to remove proteins, benzonase for the depletion of nucleic acids (RNA, DNA), and ethanol to extract lipids was added. Samples along with HA standards were then subjected for polyacrylamide electrophoresis. The tissue-produced HA was visualized on the gel by staining with “Stains All” dye (Sigma-Aldrich).

### Immunofluorescent microscopy and flow cytometry

Immunofluorescent staining and flow cytometric analysis was performed as follows. To block Fc receptors, 10^6^ cells were incubated for 15 min at 4°C with anti-CD16/CD32 mAbs. Cells were then incubated for 30 min on ice in 50 μl of PBS with 1 μg of relevant fluorochrome-conjugated or matched isotype control antibodies. The expression of PD-L1 and other markers was assessed using the fluorochrome-conjugated monoclonal antibodies from Biolegend. Cytometry data were acquired with a FACS Calibur flow cytometer (BD Biosciences). Immunofluorescence was evaluated using EVOS fluorescent imaging microscope (Life Technology).

### Time-lapse video

Tissue slices were seeded in a 24-well glass-bottom Grenier plate with 500ul of media. The surrounding wells were filled with 1000 ul of sterile PBS to provide adequate humidification. Slices were incubated for 2h before imaging. Plates were transferred to the BioTek Lionhart FX that was preheated to 37C with 5% CO2. Using 20x magnification, 5 to 10 beacons were chosen per well. To compensate for good variation, z-stacks of 15 slices of 4.2uM thickness. Images were taken at 20 min intervals for 96 h. Images processing and video rendering were done using Gen5 Image Prime 3.10 (BioTek Instruments).

### Cell isolation

Gr-1-positive cells were isolated from the spleen of MBT2-bearing by using positive selection and biotin-conjugated anti-Gr-1 antibody and MACS MicroBeads against biotin, according to the manufacturer’s instruction (Miltenyi Biotec, Auburn, CA). Briefly, splenic cells were incubated with biotin-conjugated anti-Gr-1 Abs for 15 min. After washing with cold MACS buffer stained cells were incubated with anti-biotin-microbeads (Miltenyi Biotec, Auburn, CA) for an additional 15 min and subsequently subjected to positive selection of Gr-1^+^ cells on a MACS LS column according to the manufacturer’s instructions (Miltenyi Biotec).

### Quantitative real-time PCR

Total cellular RNA was isolated with direct-zol RNA MiniPrep (Zymo Research) according to the manufacturer’s instructions. Quantitative real-time PCR analysis was performed using cDNA-specific TaqMan®Gene expression assays for murine Hyal1 and Hyal3 from Applied Biosystems were used in the study. A mouse Eukaryote 18s rRNA (assay ID: 4319413E) was used as an endogenous control.

### Human studies

Tumor tissue from 10 patients diagnosed with bladder urothelial carcinomas were collected during TURBT or radical cystectomy at the Department of Urology, University of Florida. Surgically removed tumor tissues were obtained from bladder cancer patients following informed consent and approval by the University of Florida institutional review board (IRB).

### Statistical analysis

The statistical significance between values was determined by the Student *t*-test. All data were expressed as the mean ± SD. Probability values ≥ 0.05 were considered non-significant. Significant values of p≤ 0.05 were expressed as an asterisk (*). The flow cytometry data shown are representative of at least three separate determinations.

## ACKNOWLEDGEMENTS

This work supported by grant #8JK05 from James & Esther King Biomedical Research Program and 1923 Fund to S.K.

