## Supplemental Figures for "Detection of PD-L1-expressing myeloid cell clusters in the hyaluronan-enriched stroma in tumor tissue and tumor-draining lymph nodes"

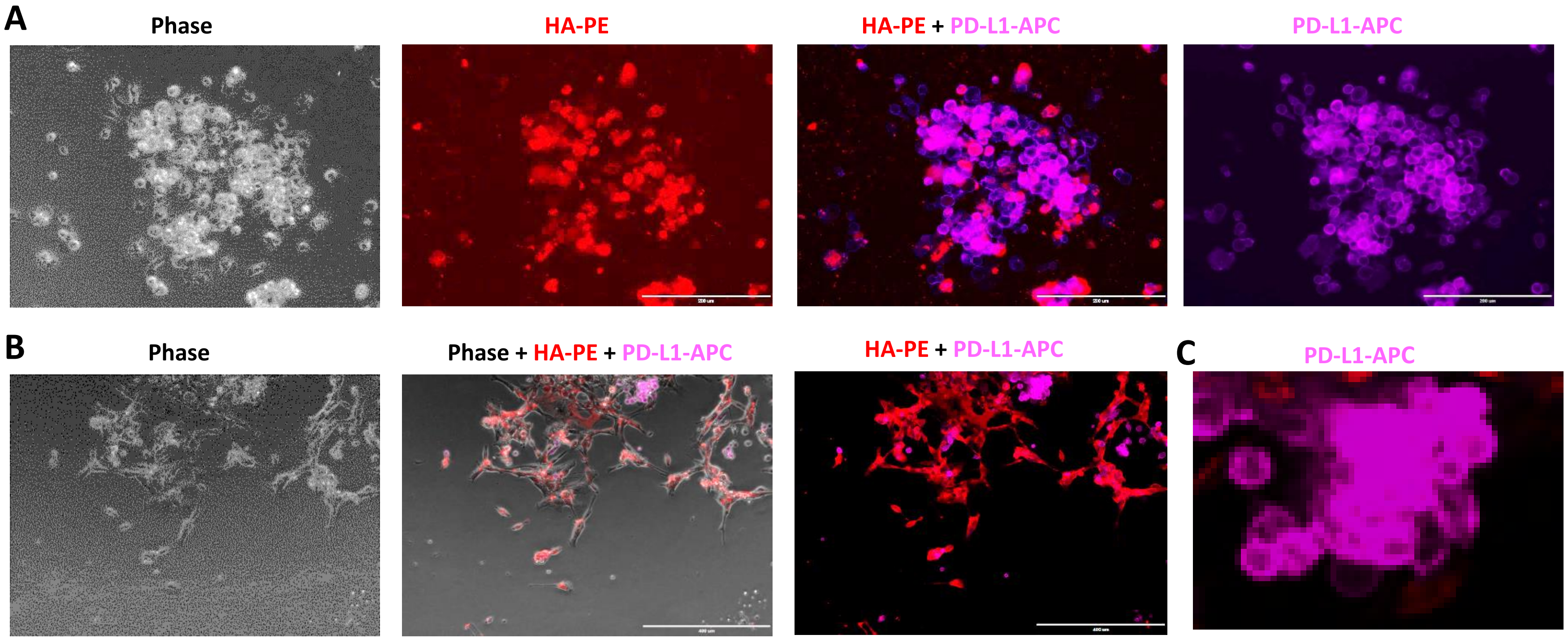


**Supporting Fig. S1. The stroma-associated HA supports development of PD-L1$^{+}$ cells in tumor tissue and tumor-draining lymph nodes.** MBT2 tumor cells were injected into C3/He mice. Two weeks later mice were sacrificed, and tumors were surgically excised. Tissue slices prepared using tumor tissue pieces (**A, B, C**). Forty eight hours later plates were fixed with 4% formaldehyde and stained for the expression of PD-L1 (magenta) and HA (red). Representative images of tumor stroma are shown.

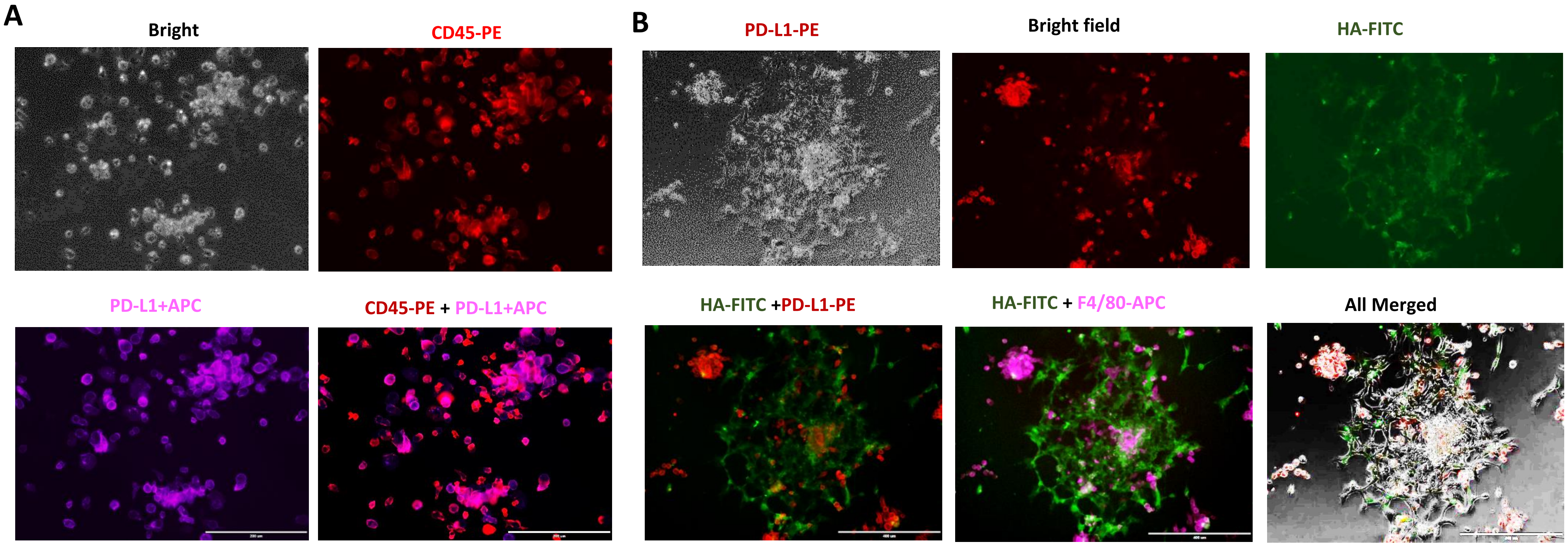


**Supporting Fig. S2. HA-enriched stroma supports development of PD-L1$^{+}$F4/80$^{+}$ macrophages**. MBT2 tumor cells were injected into C3/He mice. MBT2 tumor tissue slices were cultured in 24-well plates, allowing develop the stromal clusters and produce hyaluronan (HA). Plates were fixed with 4% formaldehyde and stained for (**A**): CD45-PE (red) and PD-L1-APC (magenta) ; and (**B**): HA (green), PD-L1 (red) and F4/80 (magenta). Representative images are shown.

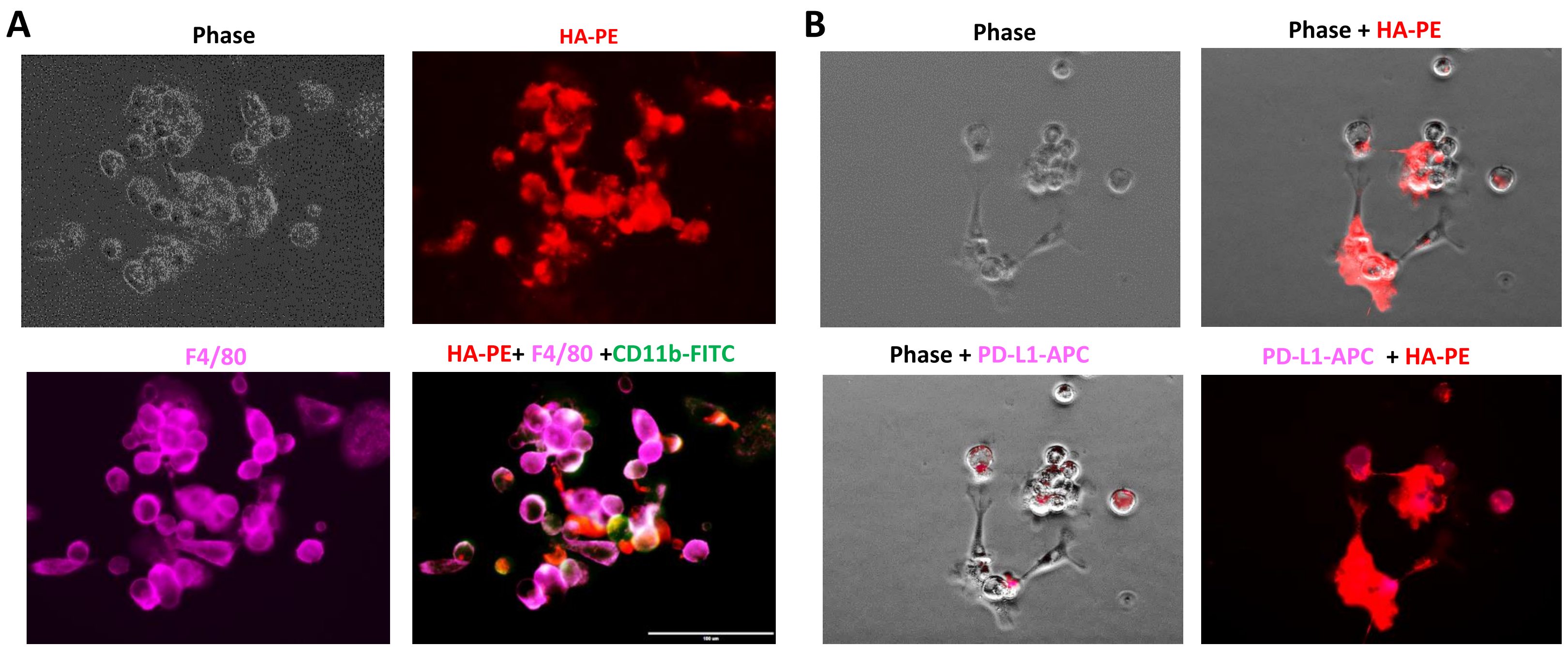


**Supporting Figure S3. PD-L1-expressing cells detected in human HA-enriched tumor stroma. A**: Murine MBT2 tumor tissue slices were cultured in 24-well plates. Plates were fixed formaldehyde and stained for HA-PE (red), F4/80 (magenta) ; and CD11b (green). Representative images are shown. **B:** Representative images showing the presence of HA-enriched stroma interacting with PD-L1+ cells in human bladder cancer tissue. Tumor tissue slices prepared cultured for 7 days. After removing culture medium and tissue slices, plates were fixed , stained for HA (red), PD-L1-APC (magenta) and evaluated by IF microscopy.

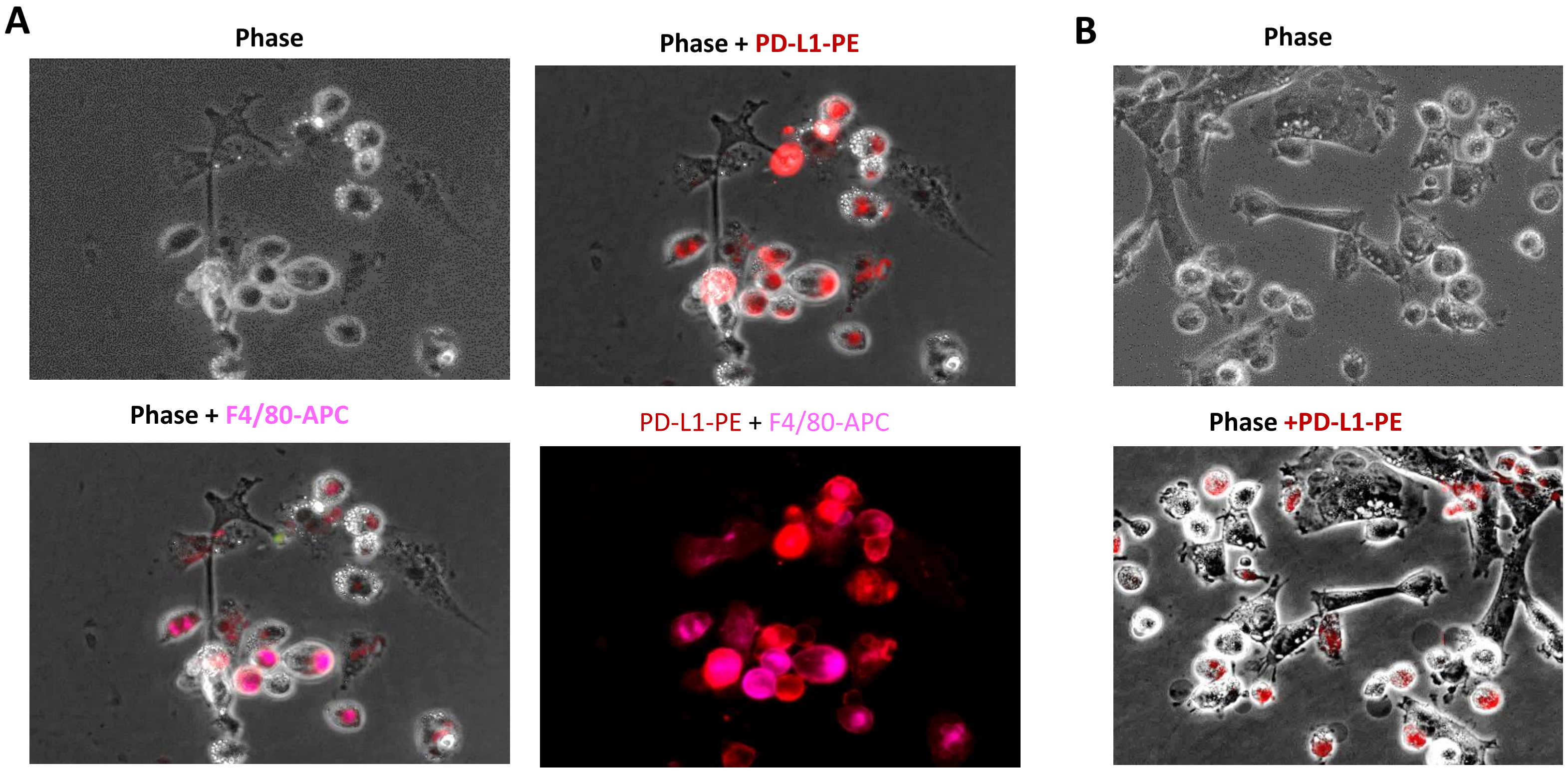


**Supporting S4. Fibroblast-like cells support development of PD-L1$^{+}$ cells in both TDLNs and tumor tissue.** MBT2 cells were injected into C3/He mice. Two weeks later, mice were sacrificed, and tumor-draining LNs (**A**) were surgically excised. Prepared tissue slices were cultured for 5 days to produce HA and attach to the plates. Plates were fixed and stained for the presence of PD-L1-PE (red) and F4/80-APC (magenta). Tumor-tissue slices (B) were analyzed by live imaging (not fixed). Anti-PD-L1-PE antibody (red) to the tissue slice cultures and pictures taken 10 minutes later. Representative images are shown.

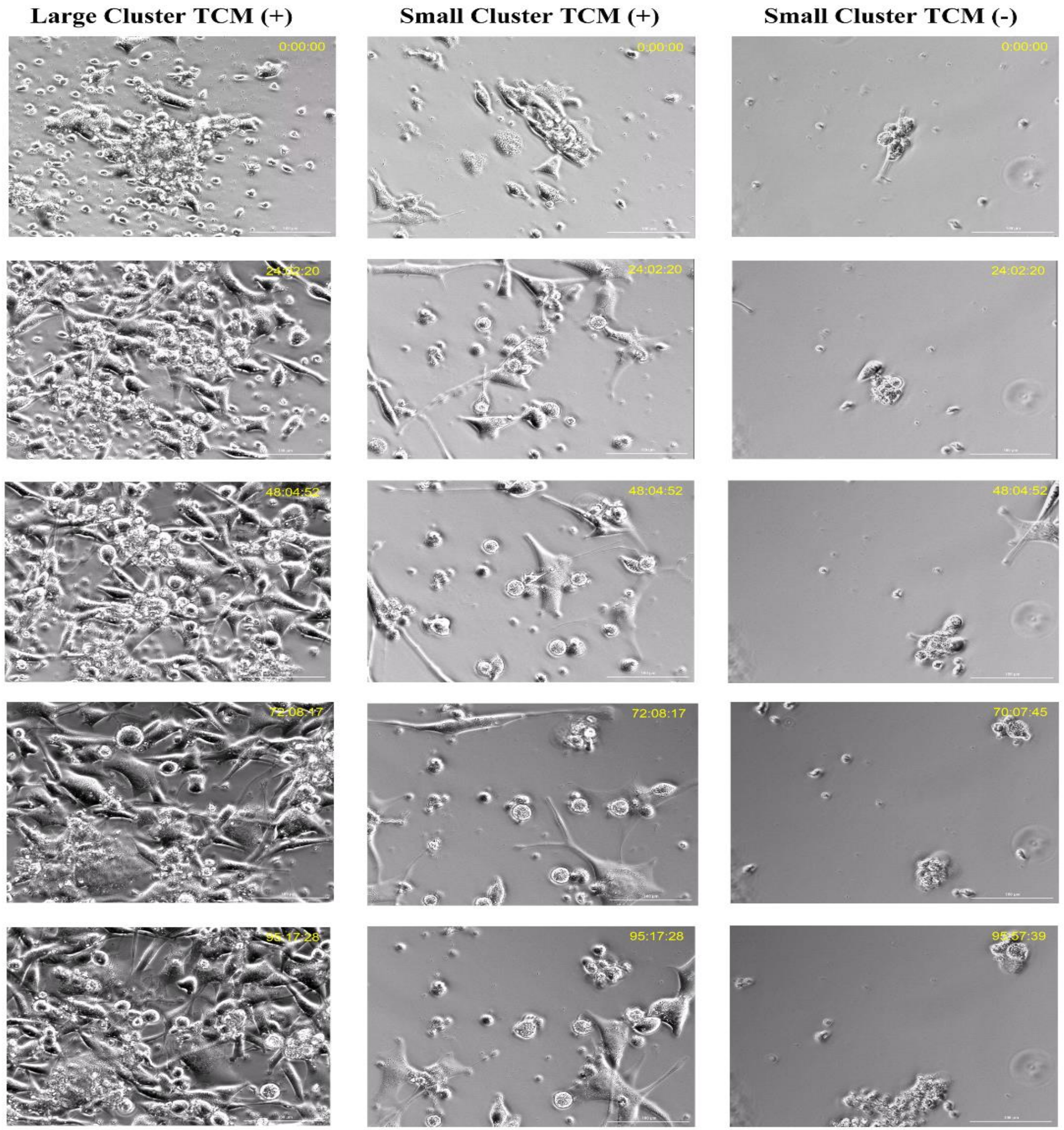


**Supporting Figure S5**. **Formation of stromal clusters using time-lapse.** Infiltrating cells from bladder cancer slice culture were imaged for 96h the presence of TCM (+) or without TCM(-). Both Large and Small Clusters TCM (+) displayed dynamic motility, proliferation, and viability through the experiment. The TCM (-) only developed small clusters that displayed some motility and proliferation; however they began to display decreased viability 72h. TCM provides factors that enhance infiltrate survival, proliferation, and motility.

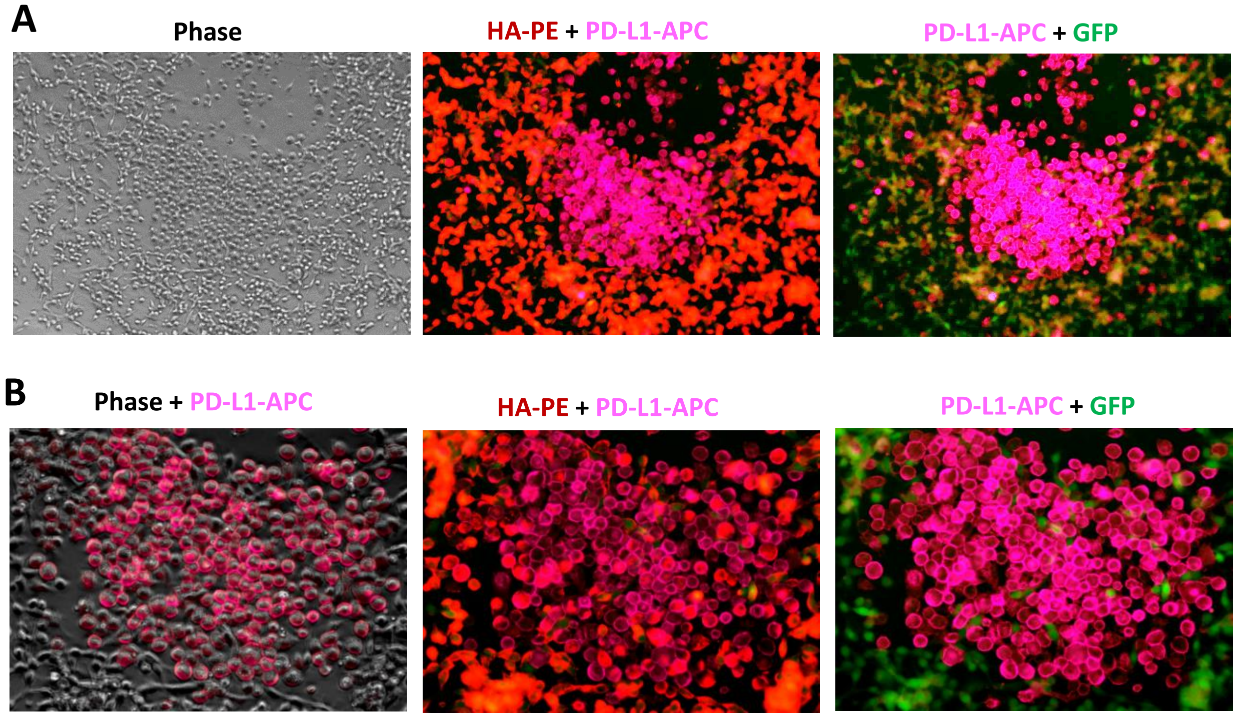


**Supporting Fig S6. A: Clusters of PD-L1$^{+}$ cells are developing in close contact with stroma that enriched with HA produced by both MBT2 tumor cells and CAFs.** GFP-expressing MBT2 (**A**) and CT26 (**B**) murune tumor cells were injected in C3/He and BALB/c mice, respectively. Two weeks later the harvested tumor was used for the preparation of tumor tissue slice culture. Tumor tissue slices were cultured for 7 days, then fixed and stained with PD-L1-APC and HA-PE Abs.

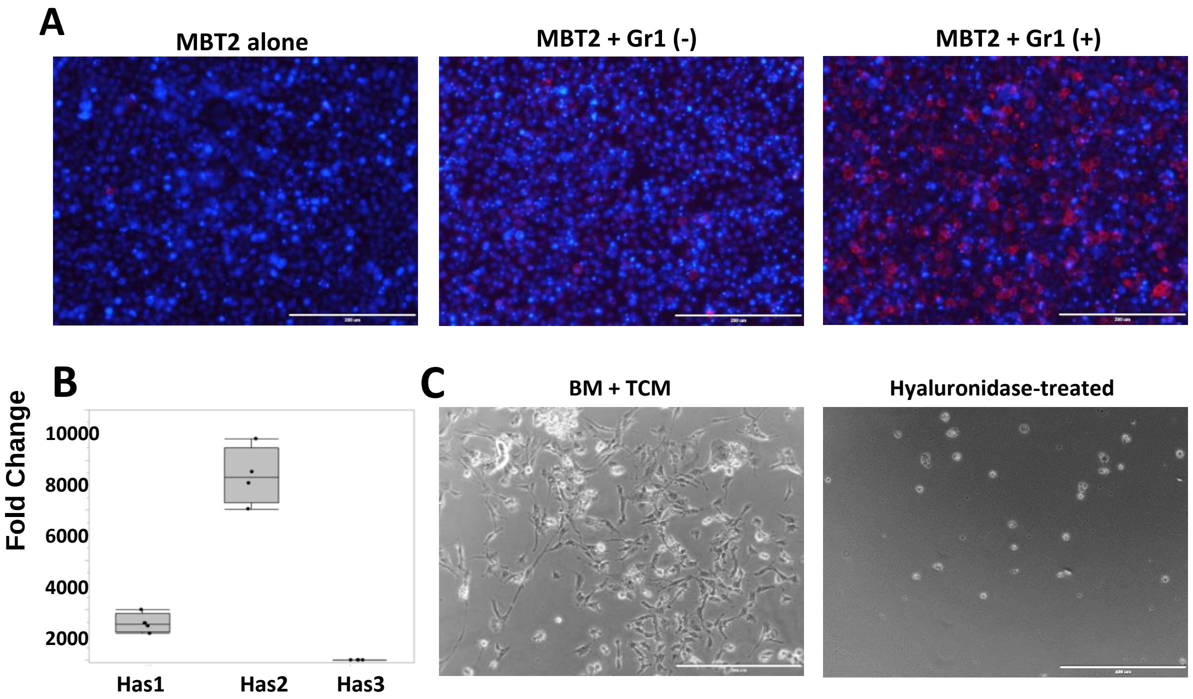


**Supporting Figure $7**: **A**: Gr-1+ cells were enriched using magnetic beads from the spleen of MBT2 bearing mice. Isolated Gr-1-positive (right panel) and Gr-1-negative (middle panel) splenic cells were mixed with MBT-2 tumor cells and added to the 24-well plate in a complete culture medium. On Day 7 cells were collected and stained with Dapi (blue) and anti-PD-L1-PE Abs (red). **B**: Expression of HAS1, HAS2 and HAS3 in MBT-2 murine bladder tumor cells. Frozen MBT2 cells were thawed and cultured for 48 hours. Collected cells were subjected to RNA isolation. Expression of murine hyaloronan synthases 1, 2 and 3 (HAS1, 2, 3) was measured using Quantitative Real-Time PCR.C: MBT2 tumo-conditioned medium added to the 24-well plate overnight with or without hyaluronidase I (Sigma-Aldrich). Myeloid cells isolated from murine bone marrow cells were added to the TCM-treated cell. Microphotographs were taken on day 7 after initioation of cell culture. Representative picures are shown.

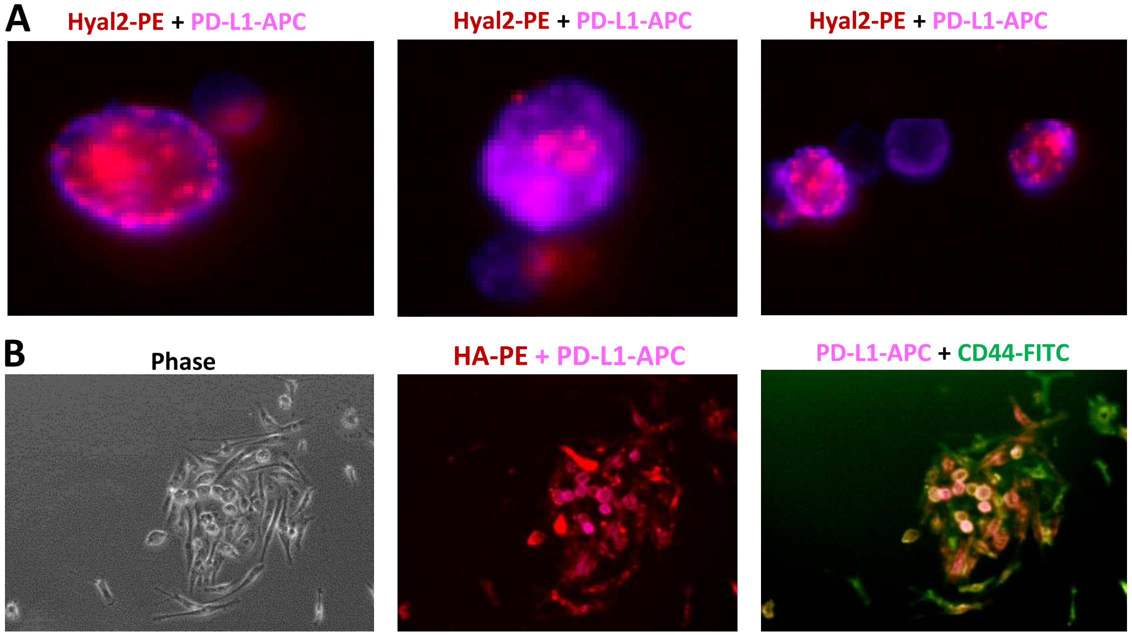


**Supporting Fig. S8. A:** CD11b myeloid cells isolated from murine naïve BM were cultured in the presence or absence of TCM. On Day 4 later cultured cells were collected. Collected cells were stained for Hyal2 (red) and PD-L1 (magenta) using immunofluorescent microscopy. **B:** Renca tumor cells were injected into BALB/c mice. Two weeks later mice were sacrificed, and tumors were surgically excised. Prepared tissue slices were cultured in the complete culture medium, fixed and stained for the HA (red), PD-L1 (magenta), and CD44 (green).

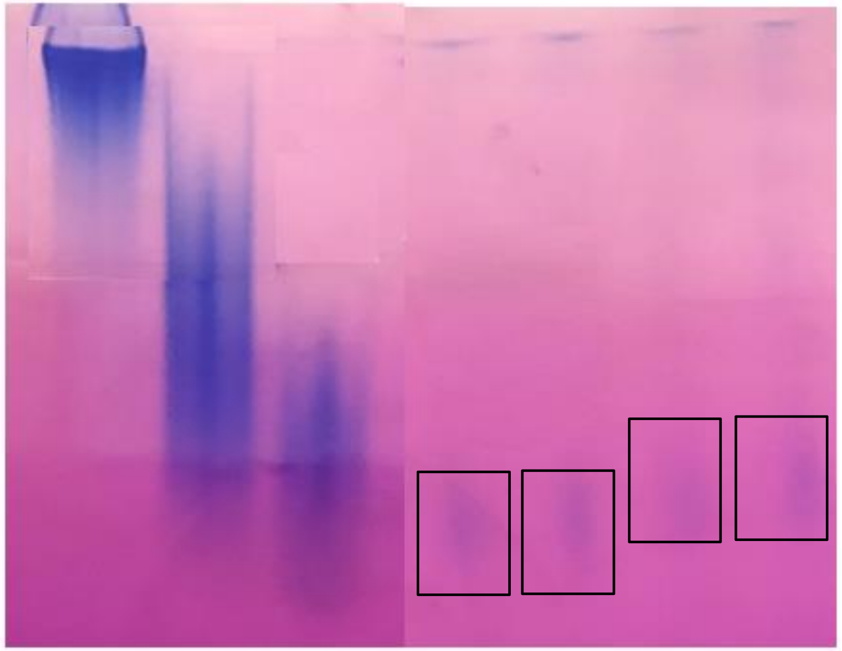


**Supporting Figure S9.** Electrophoretic analysis of HA produced by TDLNs and MBT2 tumor tissue slices. Cell-free supernatants collected from TDLN and tumor tissue slice cultures were treated with ethanol, proteinase K, and benzonase before applying samples to polyacrylamide electrophoresis**.** Commercial HA with MW 200, 10, and 5kDa were used as control. Hyaluronan was visualized by staining with "Stains All" dye.

**Supporting Fig. 9**
